# Rapid adaptive evolution of scale-eating kinematics to a novel ecological niche

**DOI:** 10.1101/648451

**Authors:** Michelle E. St. John, Roi Holzman, Christopher H. Martin

## Abstract

The origins of novel trophic specialization, in which organisms begin to exploit novel resources for the first time, may be explained by shifts in behavior such as foraging preferences or feeding kinematics. One way to investigate the behavioral mechanisms underlying ecological novelty is by comparing prey capture kinematics between groups. In this study, we investigated the contribution of kinematics to the origins of a novel ecological niche for scale-eating within a microendemic adaptive radiation of pupfishes on San Salvador Island, Bahamas. We compared prey capture kinematics across three species of pupfish while consuming shrimp and scales in the lab and found that scale-eating pupfish exhibited peak gape sizes that were twice as large as all other groups, but also attacked prey with a more obtuse angle between their lower jaw and suspensorium. We then investigated how this variation in feeding kinematics could explain scale-biting performance by measuring the surface area removed per strike from standardized gelatin cubes. We found that a combination of larger peak gape and more obtuse lower jaw and suspensorium angles resulted in 67% more surface area removed per strike, indicating that scale-eaters may reside on a performance optimum for scale-biting. We also measured feeding kinematics of F1 hybrids to test whether feeding performance could contribute to reproductive isolation between species and found that F1 hybrid kinematics and performance more closely resembled those of generalists, suggesting that they may have low fitness in the scale-eating niche. Ultimately, our results suggest that the evolution of strike kinematics in this radiation is an adaptation to the novel niche of scale-eating.

## Introduction

Determining how organisms use resources for the first time and occupy novel niches is an outstanding question in evolutionary ecology. Many changes accompany adaptation to a novel niche and previous studies have investigated how shifts in behaviors (Bowman and Billeb 1965; Tebbich et al. 2010; Curry and Anderson 2012), morphologies (Ferry-Graham et al. 2001; Ferry-Graham 2002; Hata et al. 2011; Davis et al. 2018), physiologies (Arias-Rodriguez et al. 2011; Tobler et al. 2015, 2018), and kinematics (Janovetz 2005; Patek et al. 2006; Cullen et al. 2013; McGee et al. 2013) can all facilitate this transition.

Shifts in kinematic traits—particularly those which affect prey capture and feeding—are especially promising, because they can provide biomechanical insights into the origins of novel trophic niches. For example, the wimple piranha (*Catoprion mento*) uses a ram attack coupled with a uniquely large gape angle to knock scales free from its prey (Janovetz 2005); Syngnathiform fishes specialize on evasive prey items using power-amplified jaws (Longo et al. 2018); and the Pacific leaping blenny (*Alticus arnoldorum)* is able to feed and reproduce on land by using unique axial tail twisting to improve propulsion and stability for greater jumping performance (Hsieh 2010).

Differences in prey capture kinematics between species may also contribute to postzygotic extrinsic reproductive isolation by reducing hybrid feeding performance (Higham et al. 2016b), which may lead to speciation (Henning et al. 2017; Matthews and Albertson 2017). For example, McGee et al. (2015) measured prey capture kinematics and performance in two sunfish species (Centrarchidae) and their naturally occurring hybrids. Hybrid sunfish displayed intermediate gape size compared to parental types and initiated strikes from an intermediate distance, yet their actual feeding performance was less than predicted from additive traits. Hybrid Lake Victoria cichlids (produced by crossing thick-lipped *Haplochromis chilotes* and thin-lipped *Pundamilia nyererei* parents) also exhibited lower foraging performance compared to parental species, most likely due to antagonistic pleiotropy and genetic correlations between head and lip morphology (Henning et al. 2017). Despite these findings, few studies investigate how hybrid kinematics affects the evolution of novelty or explicitly connect kinematics to performance consequences.

Scale-eating (lepidophagy) is excellent for connecting a mechanistic understanding of feeding kinematics with adaptation to a novel trophic niche. It is an extremely rare trophic niche which has independently evolved only 19 times in approximately 100 fish species out of over 35,000 (Sazima 1983; Martin and Wainwright 2013a; Kolmann et al. 2018). However, not much is known about its evolutionary origins or its kinematics. Current hypotheses for the origins of scale-eating vary, but they all take a strict behavior-first approach (Greenwood 1965; Sazima 1983; St. John et al. 2018) suggesting that kinematic variation may also contribute to the origins of scale-eating. However, only a few studies have investigated the feeding kinematics and performance of scale-eating fishes. Janovetz (2005) measured feeding kinematics of *C. mento* while consuming: 1) free floating scales, 2) whole fish, and 3) scales off the sides of fish, and found that scale-eating kinematics were different from those used during suction-feeding or biting. Interestingly, scale-eating attacks produced gape angles that ranged from 30-100% larger than those produced from consuming free-floating scales or whole fish respectively— suggesting that a larger gape is necessary for scale-eating. Furthermore, this variation in gape angle across food items was documented within individuals, indicating that scale-eating kinematics may be behaviorally mediated (Janovetz 2005). The feeding kinematics of the Lake Tanganyikan scale-eating cichlid, *Perissodus microlepis*, have also been examined (Takeuchi et al. 2012; Takeuchi and Oda 2017). While many studies of this species focus on how kinematics interacted with *P. microlepis*’ antisymmetric mouth morphology, and not on scale-eating kinematics *per se*, they have still documented a significant interaction between kinematic traits, behavior, and morphology. For example, *P. microlepis* were able to perform more successful scale-eating strikes using their dominant side (Takeuchi et al. 2012; Takeuchi and Oda 2017). A similar oral jaw antisymmetry and behavioral laterality was documented in a scale-eating characiform (*Exodon paradoxus;* Hata et al. 2011). While these studies provide valuable insights into scale-eating kinematics and performance, the lack of comparative data on the kinematics of closely related non-scale-eating species or hybrids has so far limited further investigations of the origins of scale-eating.

The overall goal of our study was to fill the following knowledge gaps and shed light on the relationship between kinematic traits and occupation of a novel niche: First, comparisons of scale-eating kinematics across scale-eating and closely related non-scale-eating outgroup species is necessary for investigating the origins of novelty. Without the comparative method it is impossible to determine which kinematic variables are unique or important for scale-eating. Second, very few kinematic studies investigate hybrid kinematics. Understanding hybrid kinematics, especially in the context of novelty, is informative because 1) impaired performance in hybrids is a form of extrinsic postzygotic isolation between species (McGee et al. 2015; Higham et al. 2016b) and 2) it can allow the decoupling of morphology, behavior, and kinematics making it easier to identify causative traits underlying performance (Holzman and Hulsey 2017). Finally, few studies connect observed variation in kinematics to variation in whole organism feeding performance (but see: Svanbäck et al. 2003; Takeuchi et al. 2012; Whitford et al. 2019). Making this connection is important because it can identify kinematic traits associated with performance tasks relevant to evolutionary fitness rather than simply describing phenotypic variation in kinematic traits, most of which may not be relevant to performance or fitness (Arnold 1983; Hu et al. 2017).

The scale-eating pupfish (*Cyprinodon desquamator*) is an excellent organism to investigate the interaction of kinematics and novelty for several reasons. First, the scale-eating pupfish evolved within a recent sympatric radiation of pupfishes on San Salvador Island, Bahamas. This radiation is endemic to a few hypersaline lakes on the island (Martin and Wainwright 2013a), which were most likely dry during the last glacial maximum 10-15 kya (Hagey and Mylroie 1995). Second, the radiation provides closely related sister taxa for kinematic comparison including: 1) the scale-eating pupfish, 2) a generalist pupfish (*C. variegatus*), and 3) a snail-eating pupfish (*C. brontotheroides*). Phylogenetic evidence suggests that scale-eating pupfish form a clade across all lakes where they are found on San Salvador and that this clade is sister to a clade containing generalists and snail-eaters (Martin and Feinstein 2014; Lencer et al. 2017), although gene flow is still ongoing among all three species (Richards and Martin 2017). All three pupfish species can be crossed in the lab to measure the kinematics and performance of hybrid phenotypes.

The morphological similarities and differences between San Salvador pupfishes have also previously been described. Specifically, 1) all pupfish species exhibit vestigial ascending processes of the premaxilla allowing for independent movement of the upper and lower jaws during jaw protrusion (Hernandez et al. 2009, 2018), and 2) scale-eating pupfish have two-fold larger, supra-terminal oral jaws compared to the smaller, terminal jaws of the generalist or snail-eating pupfish (Martin and Wainwright 2011, 2013a; Martin 2016). Their divergent morphology, along with the findings of Janovetz (2005), leads us to predict that scale-eating pupfish should have larger gapes during scale-eating attacks compared to other species of pupfish, and that this peak gape should be the result of a larger angle between the anterior tip of the premaxilla, the quadrate-articular joint, and the anterior tip of the dentary.

We investigated the interaction between kinematics and novelty in pupfishes by recording the high-speed feeding strikes of San Salvador generalist, snail-eating, and scale-eating pupfishes, along with F1 hybrids. We asked: 1) if scale-eating pupfish differed in their feeding kinematics compared to other groups during scale-eating and suction-feeding strikes, 2) whether variation in kinematics was associated with bite performance, and 3) if F1 hybrid feeding kinematics differed from parental species. Ultimately, we found that the feeding kinematics of scale-eating pupfish diverged from all other species and were not solely due to their increased oral jaw size. Instead, scale-eaters may be behaviorally mediating their feeding kinematics to optimize the surface area removed per strike, suggesting that scale-eater kinematics are a recent adaptation to scale-eating.

## Methods

### Collection and Husbandry

We used seine nets to collect generalist, snail-eating, and scale-eating pupfishes from Crescent Pond, Little Lake, and Osprey Lake on San Salvador Island, Bahamas in July, 2017 and March, 2018. Wild-caught fish were maintained in 37-75L mixed-sex stock tanks at a salinity of 5-10 ppt and temperature of 23-27°C. While in stock tanks, fish were fed a diet of frozen bloodworms, mysis shrimp, and commercial pellet foods daily. In the lab, we crossed generalist and scale-eating pupfishes from both Little Lake and Crescent Pond to produce F1 hybrid offspring. Prior to filming, pupfishes were isolated in 2L tanks to maintain individual IDs throughout the study.

### Feeding kinematics

We recorded pupfishes feeding on three different food items: frozen mysis shrimp (Mysida, Hikari Inc.), scales, and standardized gelatin cubes (dimensions: 1.5 cm x5cm X 1.5 cm × 1.5 cm cube; Repashy Superfoods, Community Plus Omnivore Gel Premix; prepared following manufacturer’s instructions). We measured feeding kinematics while fish consumed both shrimp and scales because it allowed us to ask if 1) scale-eating pupfish differed in their feeding kinematics compared to other groups, 2) if the kinematics of scale-eating strikes differ from those used during suction feeding (e.g. shrimp), and 3) if F1 hybrid feeding kinematics differed from parental species. We explicitly examined F1 hybrid kinematics in this study because lowered hybrid feeding performance may contribute to reproductive isolation between species and may shed light on the underlying genetics of feeding kinematics. We additionally measured feeding kinematics across all groups while fish consumed gelatin cubes to ask whether variation in kinematic traits affected feeding performance (i.e. bite size).

In the lab, fish freely consumed mysis shrimp, but we had to train all species to feed on scales from the sides of euthanized zebrafish (*Danio rerio*; stored frozen) and to feed from gelatin cubes (stored at 4□). For training, we isolated each fish in a 2-liter plastic tank and presented a given food item (either euthanized zebrafish or gelatin cube) daily. If a pupfish began feeding on the item, it was left in the tank until the pupfish stopped feeding. If a pupfish did not begin feeding within one minute, the food item was removed from the tank. Any pupfish that did not feed received a supplemental feeding of commercial pellet food (New Life Spectrum Thera-A, medium sinking pellets). If an individual did not feed on a training item for more than two days, we reduced supplemental feedings to once every two days to ensure that the fish was sufficiently motivated. Once pupfish reliably began feeding on either scales or gelatin cubes, we proceeded to film their feeding behaviors according to the filming protocol below. Fish were never trained on more than one item at a time, and we instead ensured that all filming was completed for a single food item before proceeding to train for the next item.

For all three food items, we used a Sony Cyber-shot DSC-RX10 III (480fps) or Sony Cyber-shot DSC-RX100 IV 20.1 MP (480fps) for high-speed video of foraging strikes. Illumination was provided by a dimmable bi-color 480 LED light (Neewer) positioned ∼0.3 m from the filming tank. Pupfish were allowed to acclimate to the lighting before feeding commenced. Fish were considered acclimated when they moved around their tank freely (usually after ∼5 minutes). For scale-eating we used forceps to hold a euthanized zebrafish horizontally in the water column and perpendicular to the front of an individual. For mysis shrimp and gelatin cubes, we dropped the food item a few inches in front of an individual. All videos were taken from a lateral perspective. Once filming for one food item was completed, the process was repeated until we filmed each individual consuming all three food items.

### Kinematic analyses

Videos were converted to image stacks and analyzed using the image processing software (FIJI; Schindelin et al. 2012). To quantify feeding performance, we measured 10 kinematic trait metrics including 1) Peak jaw protrusion, defined as the distance (mm) from the center of the orbit to the anterior tip of the premaxilla. 2) Time to peak protrusion, defined as the time (s) from the start of an attack (defined as 20% of peak gape) to peak protrusion. 3) Peak gape, defined as the distance (mm) from the anterior tip of the premaxilla to the anterior tip of the dentary. 4) Time to peak gape, defined as the time (s) from the start of an attack at 20% of peak gape to peak gape. 5) Gape angle was the angle (degrees) produced at peak gape between the anterior tip of the premaxilla, the quadrate-articular joint, and the anterior tip of the dentary. 6) Lower jaw angle was the angle produced at peak gape between the lower jaw, the quadrate-articular joint, and the ventral surface of the fish beneath the suspensorium (Figure 1&2). 7) Time to impact was the time (s) from the start of an attack (20% peak gape) to first contact of oral jaws with the prey item. 8) Time from peak gape to impact was the difference between the time to impact (s) and the time to peak gape (s). 9) Starting distance from prey was the distance (mm) from the center of the orbit at the start of an attack to the center of the orbit at impact with prey item. Finally, 10) ram speed was the starting distance from prey at 20% of peak gape (m) divided by the time to impact (s). In addition to our kinematic metrics, we also measured body length and lower jaw length (Table S1) using images from the video. We calibrated each video using a grid, positioned at the back of the filming tank.

**Figure 1.**
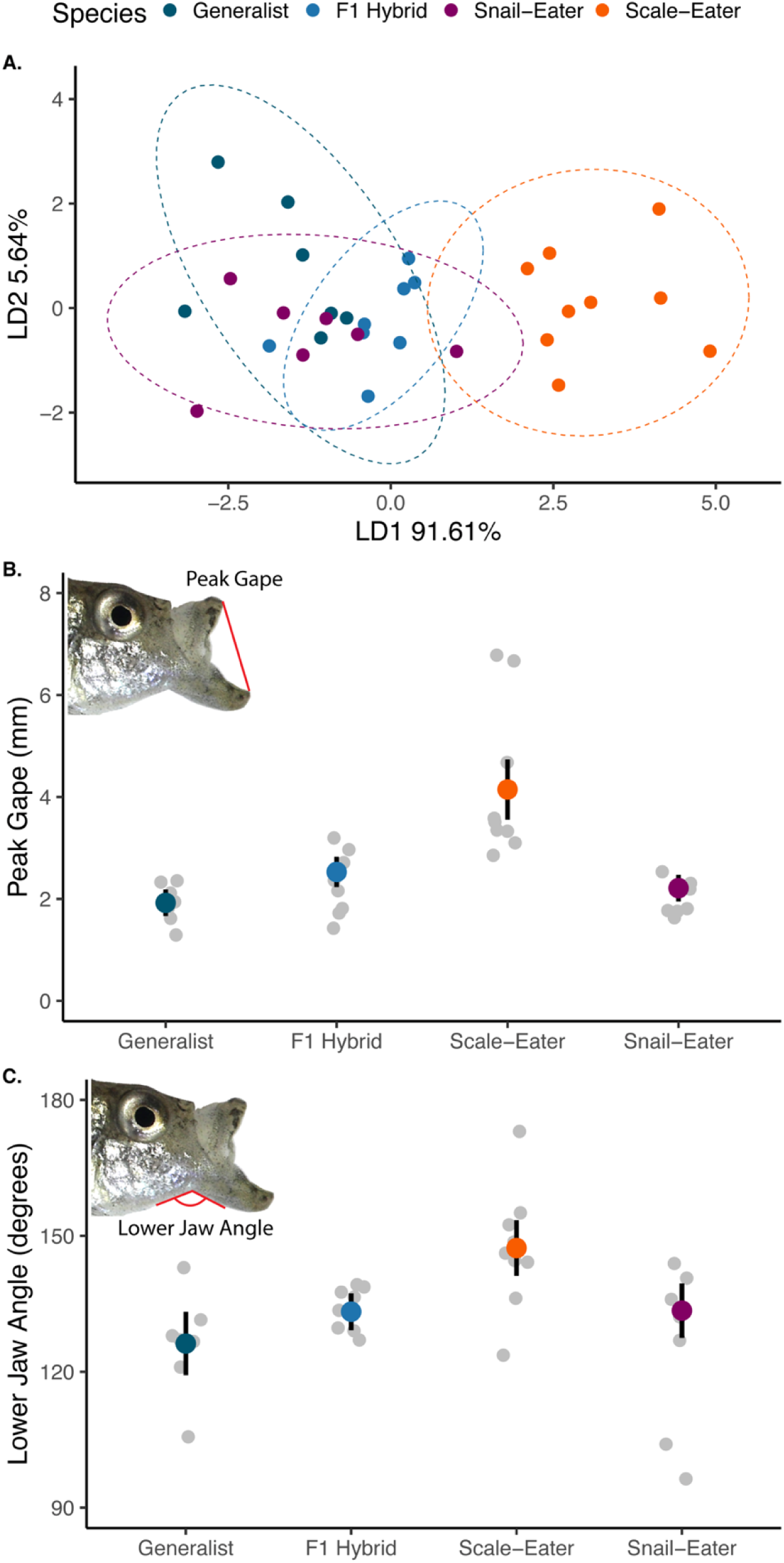
Divergent feeding kinematics in scale-eaters. A) Biplot of discriminant axes 1 (LD1) and 2 (LD2) describing overall kinematic differences among pupfish species (generalists, snail-eaters, scale-eaters, or F1 hybrids). Ellipses represent 95% CIs. B) Mean peak gape (mm) for each species with ± 95% CIs calculated via bootstrapping (10,000 iterations). C) Mean lower jaw angle at peak gape (mm) for each species with ± 95% CIs calculated via bootstrapping (10,000 iterations).

### Measuring bite performance

In order to connect variation in feeding kinematics to variation in bite size we recorded high-speed strikes on gelatin meal replacement for fish in the shape of a 1.5 × 1.5 × 1.5 cm cube. We filmed a feeding strike on a single cube and immediately removed the cube from the tank. The gel cube retains its shape in water and therefore allowed us to precisely photograph and measure the area removed by each bite. We used an Olympus Tough TG-5 camera to take photos of each lateral surface of the cube –ensuring that we had photographed the entire bite—and measured the total surface area removed from the cube (Figure 3D). One caveat is we did not measure the depth of the bite, which may be affected by additional kinematic variables during the strike. However, scale-eating attacks observed in the lab and field do not typically produce deep wounds where bite depth may be relevant, thus we expect that surface area is the best proxy for scale-biting performance in this system. Although bites were removed from both the lateral surface and edge of the gelatin cubes during strikes, there was no significant difference in surface area removed (*t*-test, *P* = 0.12).

### Statistical analyses

#### Comparing strike kinematics

We collected and analyzed 101 feeding strikes from 31 individuals striking both shrimp and scales (7 generalists; 7 snail-eaters; 9 scale-eaters; 8 F1 hybrids). We used linear mixed models (LMMs) in the lme4 package (Bates et al. 2014) in R (R Core Team 2018) to determine if any of our kinematic metrics varied between species or food item. In each model we included: 1) the kinematic metric as the response variable, 2) species designation, food item, and their interaction as fixed effects, 3) individual fish IDs and population nested within species as random effects, and 3) log body size as a covariate (Table 1). Although we compared kinematic data across multiple species, very few genetic variants are fixed between species (<1,000 SNPs out of 12 million) and generalists and molluscivores cluster by lake rather than by species (McGirr and Martin 2017; Richards and Martin 2017). Thus, is it appropriate to analyze species differences at these recent timescales as population-scale data using mixed model analyses of independent populations (e.g. Hatfield and Schluter 1999; McGee et al. 2013), rather than phylogenetic comparative methods.

**Table 1.** Results of linear mixed models investigating if strike kinematic variables vary between 1) species (generalists, snail-eaters, scale-eaters, or hybrids), 2) food item (shrimp or scales), or 3) the interaction between the two. Significant predictors are indicted in bold.

| **Response** | **Predictors** | **$\chi^2$** | ***df*** | ***P*** |
| --- | --- | --- | --- | --- |
| Peak Protrusion (mm) | Species | 4.01 | 3 | 0.26 |
|  | Food Item | 1.10 | 1 | 0.29 |
|  | log(Body Length) | 3.01 | 1 | 0.082 |
|  | Species:Food Item | 2.03 | 3 | 0.57 |
| Time to Peak Protrusion (s) | Species | 3.80 | 3 | 0.27 |
|  | Food Item | 0.73 | 1 | 0.39 |
|  | log(Body Length) | 1.02 | 1 | 0.31 |
|  | Species:Food Item | 4.03 | 3 | 0.26 |
| Peak Gape (mm) | **Species** | **23.13** | **3** | **3.8x10<sup>-5</sup>** |
|  | Food Item | 0.71 | 1 | 0.40 |
|  | log(Body Length) | 1.24 | 1 | 0.27 |
|  | Species:Food Item | 0.65 | 3 | 0.88 |
| Time to Peak Gape (s) | Species | 2.43 | 3 | 0.49 |
|  | Food Item | 0.57 | 1 | 0.45 |
|  | log(Body Length) | 2.80 | 1 | 0.17 |
|  | Species:Food Item | 1.87 | 3 | 0.60 |
| Gape Angle (degrees) | Species | 3.28 | 3 | 0.35 |
|  | Food Item | 0.032 | 1 | 0.86 |
|  | log(Body Length) | 1.01 | 1 | 0.32 |
|  | Species:Food Item | 3.43 | 3 | 0.33 |
| Lower Jaw Angle (degrees) | **Species** | **18.62** | **3** | **0.00033** |
|  | Food Item | 0.0031 | 1 | 0.96 |
|  | log(Body Length) | 3.53 | 1 | 0.060 |
|  | Species:Food Item | 3.56 | 3 | 0.31 |
| Time to Impact (s) | Species | 2.55 | 3 | 0.47 |
|  | Food Item | 2.05 | 1 | 0.15 |
|  | log(Body Length) | 1.40 | 1 | 0.24 |
|  | Species:Food Item | 4.69 | 3 | 0.20 |
| Time from Peak Gape to Impact (s) | Species | 2.44 | 3 | 0.48 |
|  | Food Item | 0.97 | 1 | 0.32 |
|  | log(Body Length) | 0.57 | 1 | 0.45 |
|  | Species:Food Item | 1.39 | 3 | 0.71 |
| Starting Distance from<br>prey (mm) |  |  |  |  |
|  | Species | 0.43 | 3 | 0.93 |
|  | Food Item | 1.99 | 1 | 0.16 |
|  | log(Body Length) | 2.77 | 1 | 0.10 |
|  | Species:Food Item | 0.80 | 3 | 0.85 |
| Ram speed (m/s) |  |  |  |  |
|  | Species | 3.25 | 3 | 0.35 |
|  | Food Item | 3.75 | 1 | 0.053 |
|  | log(Body Length) | 1.55 | 1 | 0.21 |
|  | Species:Food Item | 2.02 | 3 | 0.57 |

We also performed a linear discriminant analysis (LDA) on the combined shrimp and scales kinematic data to reduce dimensionality and identify which kinematic metrics contributed most to differences between species (Table 2, Figure 1A). We used a MANOVA and Wilks’ ⍰ to assess the significance of the LDA. We did not have enough degrees of freedom to perform these analyses with all of our kinematic variables, so we excluded time to peak protrusion and time to impact as they were highly correlated with time to peak gape (Table S2, r^2^ > .85), and also excluded distance from prey as it was highly correlated with ram speed (Table S2, r^2^ = .90). Our MANOVA ultimately included 1) peak protrusion, peak gape, time to peak gape, gape angle, lower jaw angle, time from peak gape to impact, and ram speed as response variables, 2) species designation as a predictor variable, and 3) individual ID as a random effect.

**Table 2.**
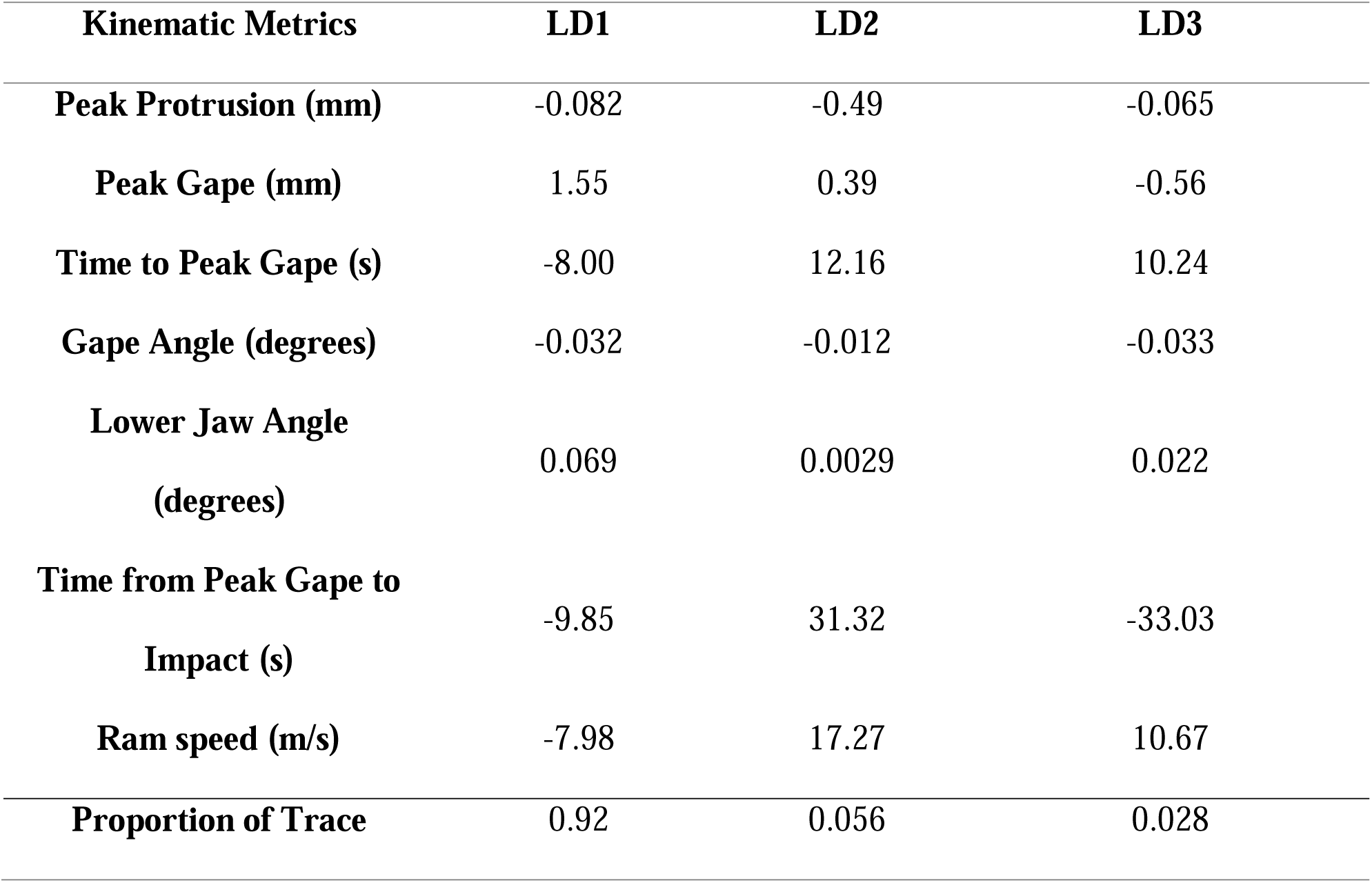
Results of a linear discriminant analysis for kinematic variables for strikes on shrimp and scales.

#### Determining how kinematic variables affect bite performance

We collected and analyzed 31 strikes on cubes across all three species and F1 hybrids. We used generalized additive models (GAMs) from the mgcv package (Wood 2011) in R to investigate how peak gape, peak protrusion, gape angle and lower jaw angle affected bite size. We used GAMs for this analysis because they do not assume a linear relationship between performance (i.e. bite size) and our given kinematic variables, but instead can fit smoothing splines to the data to test for nonlinear associations. We used AIC scores to select our optimal model (Table 3). We started with the most complete model which included 1) area removed from a cube (mm^2^) as the response variable 2) a spline modeling the interaction between two of our predictor variables, and 3) a single fixed effect. There were insufficient degrees of freedom to test all four terms at once in this model, therefore we tested all combinations of this model with our four predictor variables (Table 3A). We also tested all nested versions of this complex model by 1) removing the interaction term, but maintaining two splines and a fixed effect (Table 3B), 2) removing one spline and including three fixed effects (Table 3C), and finally 3) by testing the model with all four variables as only fixed effects (Table 3D). Ultimately, our best fitting model included area removed from a cube (mm^2^) as the response variable, univariate smoothing splines for peak gape and Gape angle, and peak protrusion as a fixed effect (average ΔAIC=24.96).

**Table 3.**
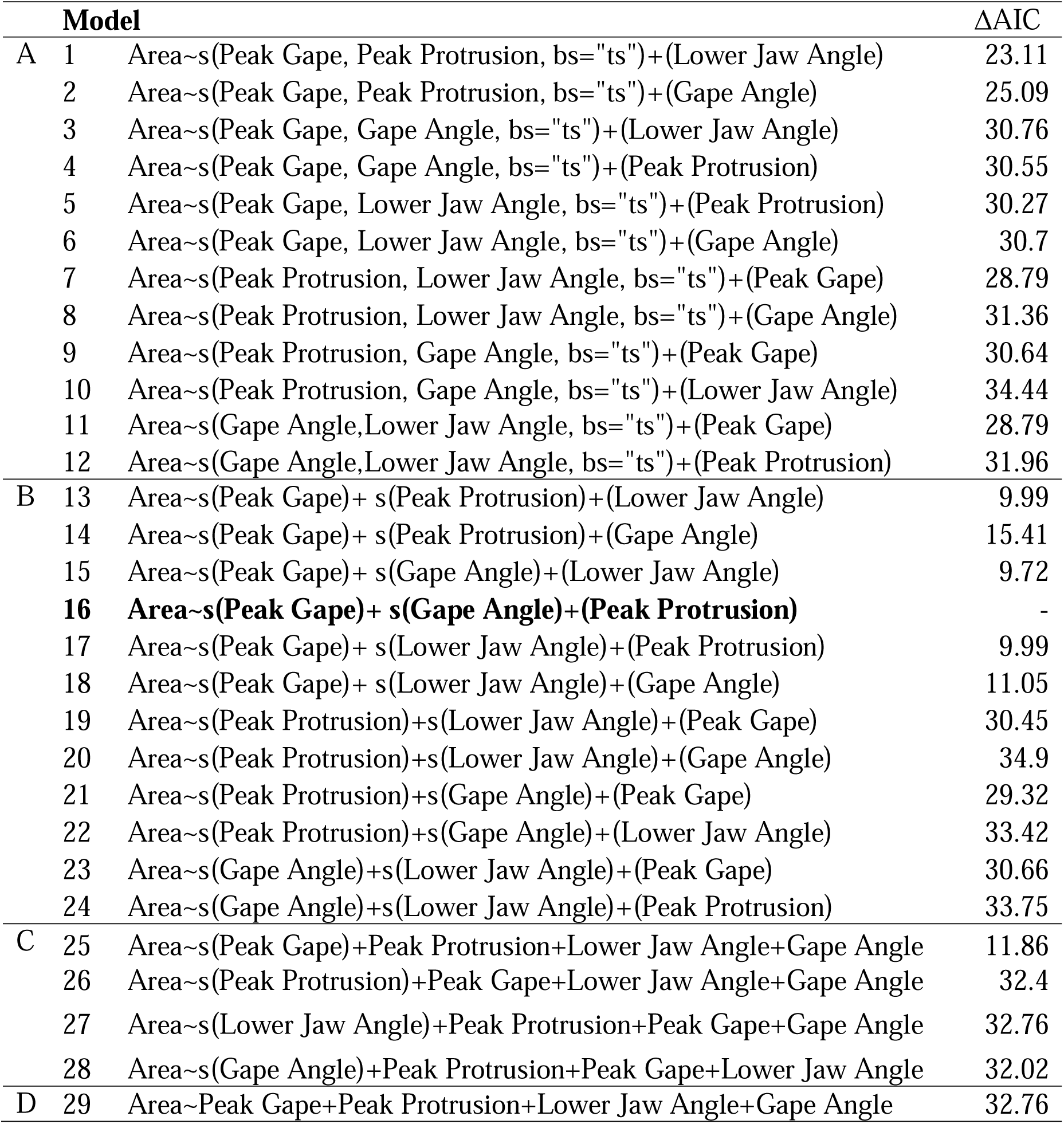
Results of GAM model comparisons using AIC score. The best-fitting model is indicated in bold and ΔAIC is indicated relative to this model.

Finally, we predicted the area removed per bite for each fish from their peak gape and gape angle kinematic measurements using a machine-learning algorithm from the caret package (Kuhn 2008) using a spline-based method. We could not measure the realized bite size for all scale-eating strikes (feeding off a euthanized zebrafish) and suction-feeding strikes (mysis shrimp). We therefore used a GAM model, estimating the effect of gape size and gape angle on the area removed from gelatin cubes, to predict bite performance (surface area removed) for all scale-biting and suction-feeding strikes in our dataset. Ideally, we would have used our best-fitting GAM model, which also included peak protrusion as a fixed effect. However, the caret package currently only accepts two fixed effects, and peak protrusion ultimately did not affect bite size (*P* = 0.078). We trained the model using all strikes observed on gelatin cubes (31 strikes across all three species and F1 hybrids) and 10-fold cross-validations with three repeats as the resampling scheme. We tested the accuracy of this model by comparing fitted values from the model to observed values from the data set and found that our model was able to predict 70% of the variance in the gelatin-strike dataset (*df*=1, F= 66.81, *P*=5.17×10^−9^, *R*^2^=0.70). We then used this model to predict the area removed for each scale-biting and suction-feeding strike based on the kinematic measurements alone. We used bootstrap resampling (10,000 iterations) to calculate mean bite size (predicted area removed) and 95% confidence intervals for each species.

#### Determining if hybrid kinematics match additive predictions

We calculated the predicted values for peak gape, lower jaw angle, and bite size for the scale-eater × generalist F1 hybrids under the hypothesis that these kinematic traits would be additive and therefore intermediate between generalist and scale-eater values. We used a one sample *t*-test to test whether the observed values of the three traits (peak gape, lower jaw angle, predicted bite sizes) for F1 hybrids deviated from additive predictions.

## Results

### Scale-eaters exhibited divergent feeding kinematics compared to other pupfishes

Scale-eaters exhibited divergent feeding kinematics, while consuming both shrimp and scales, compared to other groups (Figure 1A). A MANOVA supported the significance of this discriminant analysis and found species designation was a significant predictor of kinematics (Wilks’ ⍰ = 0.13; F = 3.05; *df* = 3; *P*= 0.00036). Species significantly varied in their peak gape and lower jaw angles during feeding strikes—regardless of the food item— in a linear mixed model controlling for individual ID and body length (Table 1). This pattern was driven by scale-eaters who had peak gapes that were twice as large as other species, but also had lower jaw angles with their suspensorium that were 14% more obtuse than other species (Figure 1B, C). Thus, the scale-eaters’ unique angle of the jaw complex with respect to their body, along with their greatly enlarged oral jaws, allows them to have increased peak gape while maintaining the same gape angle as other species (Figure 2). This may allow their upper jaws to more effectively ‘rake’ scales from the prey surface. Ram speed was the only kinematic variable that marginally varied between food items: strikes on shrimp were approximately 16% faster than those on scales (Table 1, Figure S1; *P* = 0.053).

**Figure 2.**
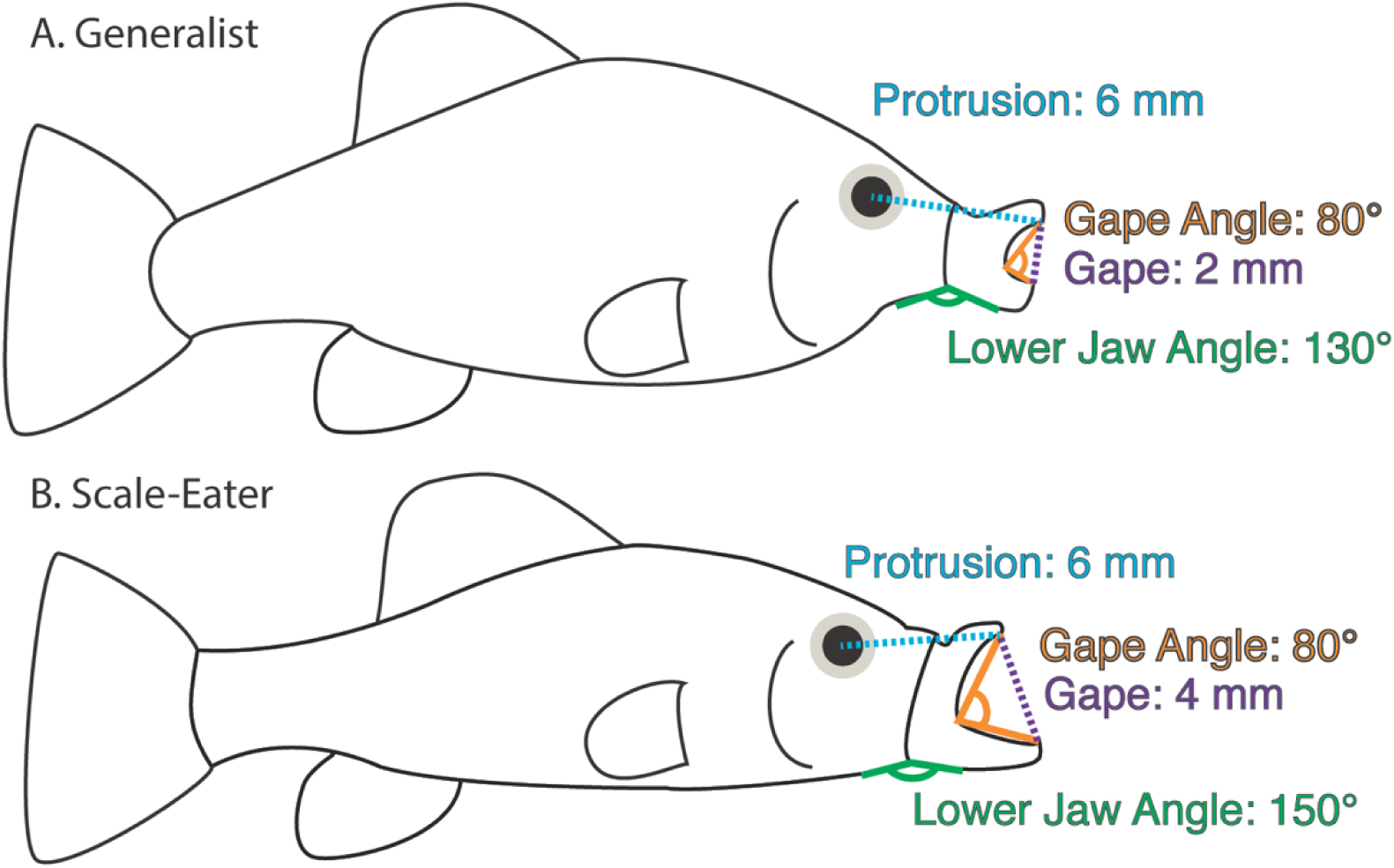
The large jaws of scale-eating pupfish allow them to double their gape size and increase the angle between their lower jaw and suspensorium (lower jaw angle) while maintaining the same gape angle as other species during feeding strikes. A) Hypothetical measurements of a generalist’s protrusion distance, peak gape, and lower jaw angle if they strike a food item with an 80° gape angle. B) Hypothetical measurements of a scale-eater’s protrusion distance, peak gape, and lower jaw angle if they strike a food item with the same 80° gape angle.

**Figure 3.**
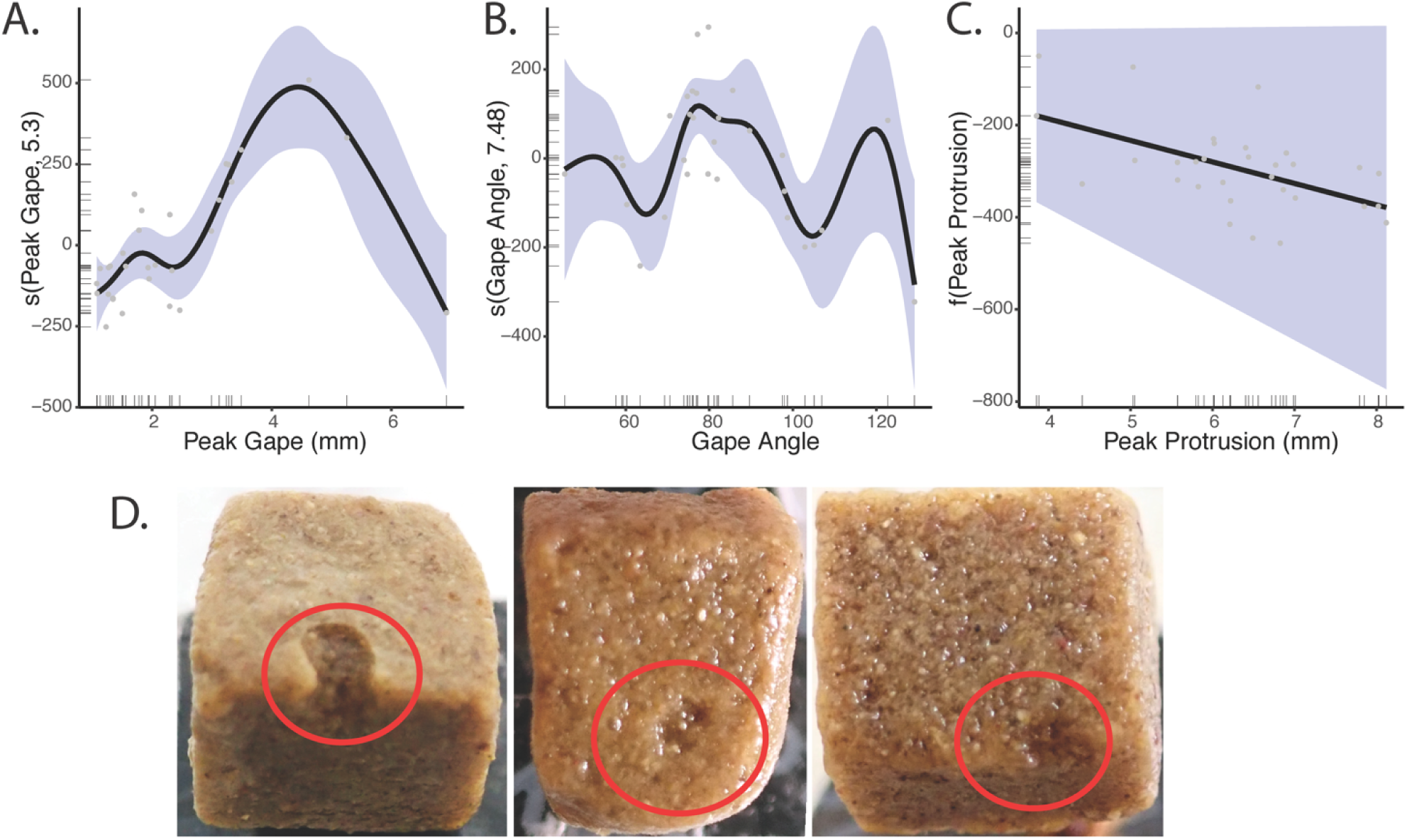
The interaction of peak gape and peak protrusion may result in a performance optimum for scale-biting. Visualization of our GAM model investigating how the thin-plate univariate splines of A) peak gape (mm), B) gape angle (degrees), and C) the fixed effect of peak protrusion (mm) is associated with bite size (surface area removed per strike from a Repashy gelatin cube), and D) representative scale-eating bites taken out of gelatin cubes.

### Variation in strike kinematics affected bite size performance

GAM modeling indicated that the univariate splines of peak gape and gape angle were both significantly associated with bite size (peak gape: *edf*=5.30, *F*=8.96, *P*= 6.33×10^−5^; gape angle: *edf* = 7.48, *F*= 2.78, *P*=0.032). The fixed effect of peak protrusion, however, was only marginally significant (*t*=−1.88, *P*=0.078). Ultimately this model explained 84.5% of the observed deviance in bite size, and suggests that a larger gapes paired with gape angles of 80° are associated with larger surface areas removed per bite (Figure 3A-C).

### F1 hybrid kinematics are not strictly additive and more closely resemble generalist kinematics

F1 hybrid feeding kinematics differed from scale-eater kinematics (Tukey HSD, peak gape: *P* = 1.2×10^−6^, lower jaw angle: *P* = 0.0090), but were not significantly different from generalist kinematics (Tukey’s HSD, peak gape: *P* = 0.21, lower jaw angle: *P* = 0.37). Mean hybrid peak gape was 39% smaller than scale-eater peak gape and 32% larger than generalist peak gape (Figure 1B). Similarly, mean hybrid lower jaw angle was 9.5% more acute than scale-eater peak lower jaw angle, and 5.6% more obtuse than the mean generalist lower jaw angle (Figure 1C). F1 hybrids failed to match additive predictions of intermediate kinematics for peak gape (t-test, μ= 3.035, mean= 2.52 mm, *P* = 0.013), but did meet these predictions for lower jaw angle (t-test, μ= 136.5, mean= 133.92 degrees, *P = 0*.*18*). Our machine learning model also predicted that scale-eater kinematics would result in bite sizes that are 60% larger than the predicted bites of the other species (Figure 4). Interestingly, estimates for F1 hybrid bite sizes were slightly smaller than expected based on additive predictions (*t*-test, μ= 320.5, mean= 255.24, *P =* 0.13).

**Figure 4.**
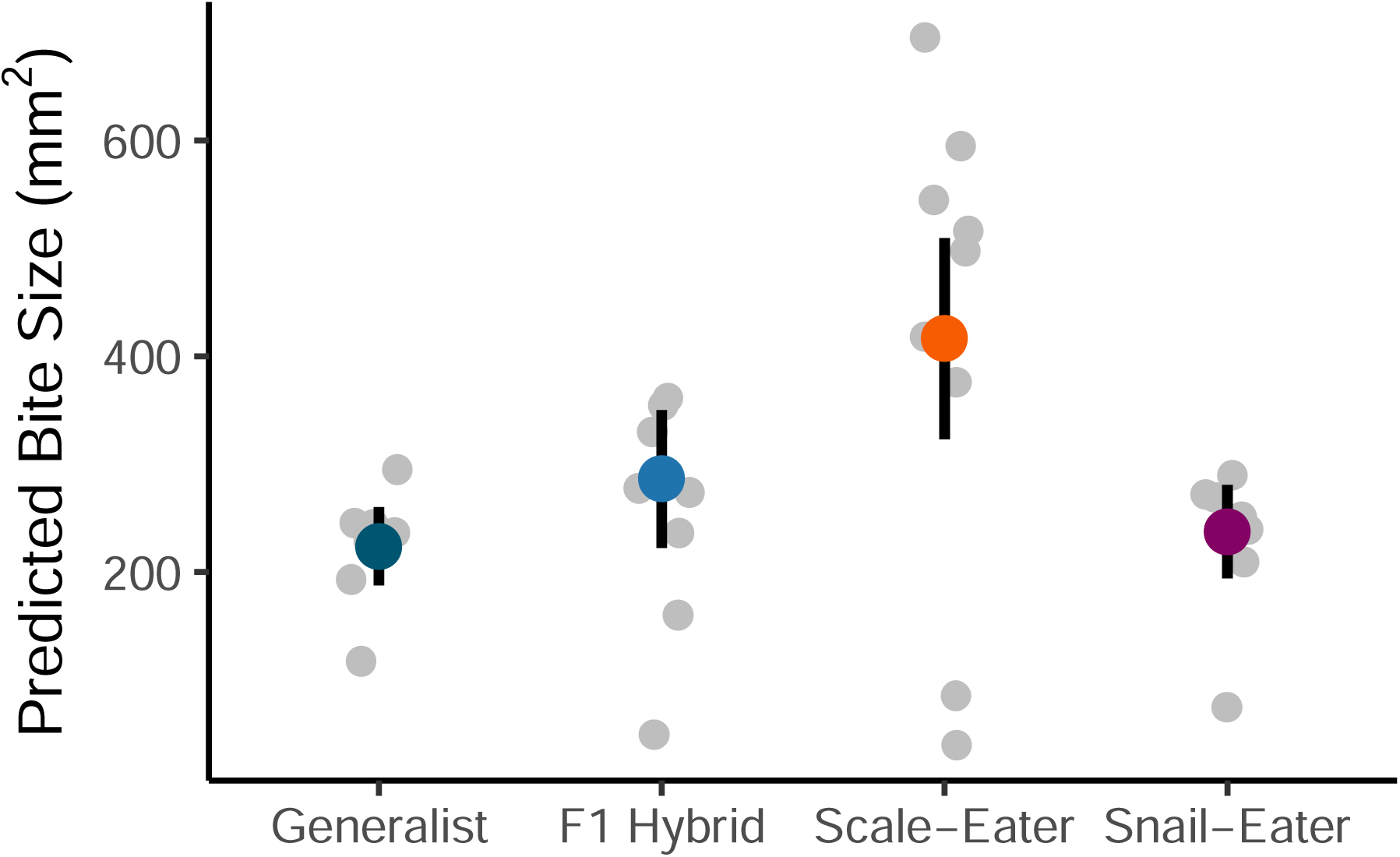
Scale-eaters have larger predicted bite sizes compared to other species. Predicted bite sizes for all strikes from each species using machine-learning optimization of GAM models. Grey points represent predicted bite sizes for individuals, color points represent means, and bars represent ± 95% CIs calculated via bootstrapping (10,000 iterations).

## Discussion

### Scale-eating pupfish have divergent feeding kinematics

Scale-eating pupfish exhibited peak gapes that were twice as large as other groups, but simultaneously displayed gap angles that were not different from other groups, and lower jaw angles that were 12% more obtuse. Thus, scale-eaters kept their jaws more closed during strikes compared to other species, resulting in smaller gape sizes than the maximum achievable gape given their morphology. These counterintuitive results only partially support our prediction that scale-eaters should have larger peak gapes, similar to the findings of Janovetz (2005) for the scale-eating piranha. Increased gape size in scale-eating pupfish was not due to an increased gape angle as we predicted. Instead, scale-eaters appear to maintain the same gape angle of their oral jaws as in other species (∼80°) and increased their lower jaw angle resulting in more closure of their jaws during strikes, effectively lowering the maximum peak gape possible due to their enlarged oral jaws (Figure 2). For example, if a scale-eater decreased their lower jaw angle from 150° to 130°, the mean lower jaw angle of generalists, they could increase their peak gape by about 8%. One possibility is that this more obtuse lower jaw angle is an artifact of filming scale-eating strikes in the lab. To investigate this, we analyzed four scale-eating strikes performed by wild scale-eaters observed in Crescent Pond, San Salvador Island, Bahamas (filmed using a Chronos camera (Kron Technologies, model 1.4, 16 GB memory, Color image sensor) with an f1.4 zoom lens in a custom underwater housing (Salty Surf, Inc. Krontech Chronos 1.4 housing with M80 flat port)) and compared their jaw angles to those seen in the lab. Wild strikes had an even more obtuse mean lower jaw angle of 168° while scale-eating strikes in the lab had a mean lower jaw angle of 153° suggesting that an obtuse lower jaw angle is also used during natural scale-eating strikes in hypersaline lakes on San Salvador Island.

Strike kinematics did not vary across prey items (Table 1), contrary to Janovetz (2005). In fact, the only kinematic variable that remotely varied between prey items was ram speed (Table 1, Figure S1), but this may simply be due to the fact that shrimp were a moving target during feeding trials while euthanized zebrafish were held stationary with forceps for scale-eating strikes. Alternatively, phenotypic plasticity due to rearing environment could produce a similar pattern; however, we did not observe any differences in strike kinematics between wild caught and lab-reared fish.

### Is jaw morphology solely responsible for kinematic variation?

The kinematic variables that varied the most between scale-eating and non-scale-eating pupfishes were peak gape and lower jaw angle—both related to the size of the oral jaws. Previous work has documented that the oral jaws of scale-eating pupfish are two-fold than their sister species (Holtmeier 2001; Martin and Wainwright 2013a; Martin 2016) and may be controlled by four moderate-effect QTL with all positive effects on jaw size, consistent with directional selection on this trait (Martin et al. 2017). It may be that increased oral jaw size is sufficient to create variation in feeding kinematics without an accompanying shift in behavior.

Previous studies have documented how changes in morphology alone can alter feeding kinematics. For example, kinematic studies have found that the scaling of the lower jaw in bluegill (Wainwright and Shaw 1999) and body size in largemouth bass (*Micropterus salmoides;* Richard and Wainwright 1995) both significantly affected prey capture kinematics. Furthermore, Ferry-Graham et al. (2010) used the pike killifish (*Belonesox belizanus*) to show that simply doubling the length of the jaws significantly affected key kinematic variables such as peak gape size—even while keeping lower jaw angle constant. Simply stated, the key adaptation necessary for scale-eating may be enlarged, supra-terminal jaws. If this hypothesis were true, we would expect that peak gape would increase with jaw size and that gape angle would increase with the shift from terminal to supra-terminal jaws, but all other kinematics variables would remain constant across species. Our results reject this hypothesis. Instead, scale-eaters maintain the gape angle observed in other species and increase their lower jaw angle with the suspensorium by 12 degrees resulting in a reduction in their potential peak gape size (Figure 2). This suggests that scale-eaters have evolved more obtuse lower jaw angles during strikes to increase feeding performance (Figures 3&4).

### Scale-eating performance optimum

Scale-eaters may have reduced their lower jaw angles relative to other species in order to remain on a performance optimum for scale-eating. Our models of bite performance supported this: peak gapes larger than approximately 4.5 mm counterintuitively resulted in smaller bite sizes (Figure 3A&B). An enlarged angle in scale-eating pupfish results in a lower jaw that points directly towards the prey during strikes – possibly resulting in greater stability for biting scales while retaining a large gape (Figure 2). This large gape and unique jaw alignment may allow scale-eaters to attack prey from a roughly perpendicular angle (as frequently observed during field observations) —appearing to wrap their large lower jaw under prey items and subsequently scraping scales from their sides using their independently protrusible upper jaws (also observed in a scale-eating characin: Hata et al. 2011). Interestingly, perpendicular angles of attack and large gapes are associated with scraping in benthic feeding fish (Van Wassenbergh et al. 2008; O’Neill and Gibb 2013). In fact, one prominent hypothesis for the origins of scale-eating is that it arose from an algae-scraping ancestor (Sazima et al. 1983). One caveat for this hypothesis, however, is that our current performance estimates do not include all possible combinations of peak gape and lower jaw angle and few observations of the largest peak gape sizes. Future work should estimate performance across multiple performance axes (e.g. Stayton 2019, Keren et al. 2018, Dickson and Pierce 2019), ideally using F2 hybrids. F2 hybrids are a useful tool for this type of experiment, as they are the first generation of offspring where recombination has the potential to produce new combinations of kinematic, morphological, and behavioral traits not observed in the F0 or F1 generations.

### Non-additive F1 hybrid feeding kinematics may contribute to reproductive isolation of scale-eaters

It is well documented that complex performance traits, such as feeding kinematics, are most likely controlled by numerous loci (i.e. polygenic), and can mostly be described as additive (reviewed in Sella and Barton 2019). We therefore expected F1 hybrids to exhibit intermediate kinematics and performance relative to both parental species. Instead, we found that F1 hybrid kinematics more closely resembled generalists (Table 1; Figure 1) and that their estimated bite sizes are slightly below additive estimates (additive prediction= 341.48, predicted = 289.72, *P = 0*.*058*) suggesting that F1 hybrids may have higher performance in a generalist trophic niche. Current evidence from field fitness experiments supports the idea that hybrid pupfish exhibit better performance in the generalist ecological niche compared to their performance in the scale-eater niche. One field experiment in these lakes measured hybrid fitness in the wild and found high mortality and low growth rates for hybrids most closely resembling the scale-eating phenotype (Martin and Wainwright 2013b). Furthermore, for the few hybrids resembling scale-eaters which did survive, only 36% had recently consumed any scales compared to 92% of wild-caught scale-eaters (Martin and Wainwright 2013a,b). Impaired hybrid performance in the scale-eating niche may contribute to extrinsic postzygotic isolation between species (McGhee et al. 2007; McGee et al. 2013; Higham et al. 2016). Reproductive isolation may also evolve more quickly in species that occupy a more distant fitness peak with a larger fitness valley such as the scale-eating pupfish due to stronger selection against hybrids and reinforced pre-mating isolation (Martin and Feinstein 2014). Thus, impaired hybrid scale-eating performance could also contribute to increased diversification rates through the mechanism of a wider fitness valley.

Low hybrid performance may also be due to the morphological differences between scale-eaters and generalists. As mentioned above, it is possible that a shift in morphology – such as enlarged oral jaws in scale-eaters—may be sufficient to change kinematic profiles alone. F1 hybrid kinematics clearly differed from scale-eater kinematics, but their jaw lengths were also significantly smaller than the jaws of scale-eaters (Tukey’s HSD, *P =* 5.21×10^−5^). Furthermore, previous work has shown that F1 hybrid pupfish offspring (produced from generalist × scale-eater crosses) tend to develop along a more similar trajectory to their maternal parent (Holtmeier 2001). This could indicate that F1 hybrid pupfish with scale-eating mothers are more likely to develop jaws resembling a purebred scale-eater, but may also retain their generalist-like kinematics. The resulting mismatch between morphology, kinematic traits, and ecological niche may be driving low hybrid survival in the scale-eating niche and contributing to reproductive isolation between generalist and scale-eating pupfish species.

## Conclusion

In conclusion, this study suggests that shifts in kinematic traits may have preceded or facilitated the origin of scale-eating in *Cyprinodon* pupfishes. Scale-eating pupfish exhibited peak gapes that were twice as large as other pupfish species, but simultaneously had lower jaw angles that were significantly larger. Surprisingly, we found that this unique combination of scale-eater kinematics may reside on a performance optimum, as large peak gapes and large lower jaw angles resulted in larger surface areas removed per strike. Impaired F1 hybrid kinematics and performance in the scale-eating niche also suggests that kinematic traits contribute to reproductive isolation of the scale-eating pupfish and the evolution of novelty. Future work should investigate if other performance optima exist on the kinematic landscape and whether F2 hybrid fitness in the wild is reduced due to a mismatch between morphology and feeding kinematics.

## Data Accessibility

All analyses reported in this article can be reproduced using the data provided by St. John and Martin (2019). Raw data will be deposited in Dryad.

## Acknowledgements

We thank the University of North Carolina at Chapel Hill, NSF CAREER 1749764, NIH 5R01DE027052-02, and BSF 2016136 for funding to CHM and the UNC Quality Enhancement Plan for course-based undergraduate research (CURE) funding to CHM. The Bahamas Environmental Science and Technology Commission and the Ministry of Agriculture kindly provided permission to export fish and conduct this research. Rochelle Hanna, Velda Knowles, Troy Day, and the Gerace Research Centre provided logistical assistance in the field; Kristi Dixon, Casey Charbonneau, Gabriel Harris, Amelia Ward, Delaney O’Connell, and the undergraduate students of BIO221L ‘Evolution of Extraordinary Adaptations’ assisted with high-speed videography and kinematic analysis. All animal care protocols were approved by the University of California, Berkeley and the University of North Carolina at Chapel Hill Animal Care and Use Committees.

